# Prioritized neural processing of social threats during perceptual decision-making

**DOI:** 10.1101/859942

**Authors:** M. El Zein, R. Mennella, M. Sequestro, E. Meaux, V. Wyart, J. Grèzes

**Affiliations:** Adaptive Rationality Center (ARC), Max-Planck for Human Development, Berlin, Germany; Cognitive and Computational Neuroscience Laboratory (LNC^2^), INSERM U960, DEC, Ecole Normale Supérieure, PSL University, 75005, Paris, France; Laboratory of the Interactions between Cognition Action and Emotion (LICAÉ, EA2931), UFR STAPS, Université Paris Nanterre, Nanterre, France; Institut du Psychotraumatisme de l’Enfant et de l’Adolescent, Conseil Départemental Yvelines et Hauts-de-Seine, Versailles, France

**Author notes:** equal contribution. **Contact**: Julie Grèzes –.

**Keywords:** perceptual decisions, EEG, facial emotions, threat, motor signals

## Abstract

Emotional signals, notably those signaling threat, benefit from prioritized processing in the human brain. Yet, it remains unclear whether perceptual decisions about the emotional, threat-related aspects of stimuli involve specific or similar neural computations compared to decisions about their non-threatening/non-emotional components. We developed a novel behavioral paradigm in which participants performed two different detection tasks (emotion vs. color) on the same, two-dimensional visual stimuli. Electroencephalographic (EEG) activity in a cluster of central electrodes reflected the amount of perceptual evidence around 100ms following stimulus onset, when the decision concerned emotion, not color. Second, participants’ choice could be predicted earlier for emotion (240ms) than for color (380ms) by the mu (10Hz) rhythm, which reflects motor preparation. Taken together, these findings indicate that perceptual decisions about threat-signaling dimensions of facial displays are associated with prioritized neural coding in action-related brain regions, supporting the motivational value of socially relevant signals.

## Introduction

Accurate decoding of socially relevant information emitted by others, such as emotional expressions, is crucial to guide adaptive decisions. Indeed, social signals are granted preferential processing: people quickly allocate attention to human faces and bodies in natural scenes ^1^, detect changes in faces better than in non-social objects ^2^, and respond faster to social than to non-social hazards ^3^. Emotional expressions, especially when they signal threat, are further prioritized relative to neutral displays and to non-social stimuli ^4–6^. Social threat signals, such as angry or fearful facial expressions, are associated with perceptual and attentional advantages ^6–11^ and they also modify the perception of surrounding environment ^12–15^. Moreover, they shape observer behavior, by increasing motor preparation ^e.g. 16–21^ and by influencing approach/avoidance decisions ^22–26^.

Perceptual decision made on non-socioemotional characteristics of stimuli show that stimulus evidence is encoded over associative, centro-parietal, regions (CPP, gradual buildup starting from 170ms after stimulus onset) and over motor regions (LRP, gradual buildup from 320ms, both CCP and LRP peak around 400-500ms ^27,28^, see also ^29–35^). Interestingly, using a perceptual decision task comparing high– and low-threat facial displays, El Zein et al. ^36^ found that high-threat displays elicited earlier (∼200ms) and enhanced stimulus encoding in both associative and motor regions simultaneously. Despite the similarity in the neural dynamics underlying perceptual decisions on threatening and neutral stimuli, the question remains whether stimulus encoding is faster over centroparietal and motor areas during perceptual decisions on socioemotional versus non-emotional stimuli.

Several methodological constraints in previous research have precluded a direct comparison between these two types of stimuli. Past studies have focused either on perceptual decisions on emotional displays ^36–39^ or on social vs non-social stimuli ^40–46^. To our knowledge, only one study has compared decisions on static facial displays of emotion and animal pictures and concluded that there is a common neural signature for emotional and non-emotional decisions ^47^. However, neither the amount of perceptual evidence nor the decision difficulty was equalized between the two tasks. Therefore, no previous study has precisely characterized the neural mechanisms involved in decisions on perceptual stimuli that simultaneously vary on threat and another non-threatening neutral dimension in a controlled, equalized way.

To reach a comprehensive understanding of whether threat expressions are prioritized during decision-making, we have developed a novel electroencephalography (EEG) paradigm in which participants performed two different detection tasks on the same, two-dimensional visual stimuli. We used facial expressions of anger as threat stimuli, signaling potentially harmful intentions. We presented morphed facial expressions (from neutral to angry) with a morphed color background (from grey to violet). In different blocks, participants were asked to report the presence or absence of either emotion (anger) or color (violet) in the stimulus, while ignoring the other task-irrelevant dimension (see **Figure 1**). Since the association of angry faces with direct and averted gaze has been shown to be appraised as high and low threat, respectively ^e.g. 48–51^, we manipulated the contextual significance of the displayed emotion via changes in gaze direction. Gaze direction was never mentioned to participants and was irrelevant to task performance. Importantly, to match the difficulty of emotion and color perceptual decisions, we not only equalized the amount of sensory evidence in the stimuli across tasks using morph matching, but also equalized the decision difficulty across the two stimulus dimensions (emotion and color) using an adaptive Bayesian titration procedure on participants’ sensitivity.

**Figure 1:**
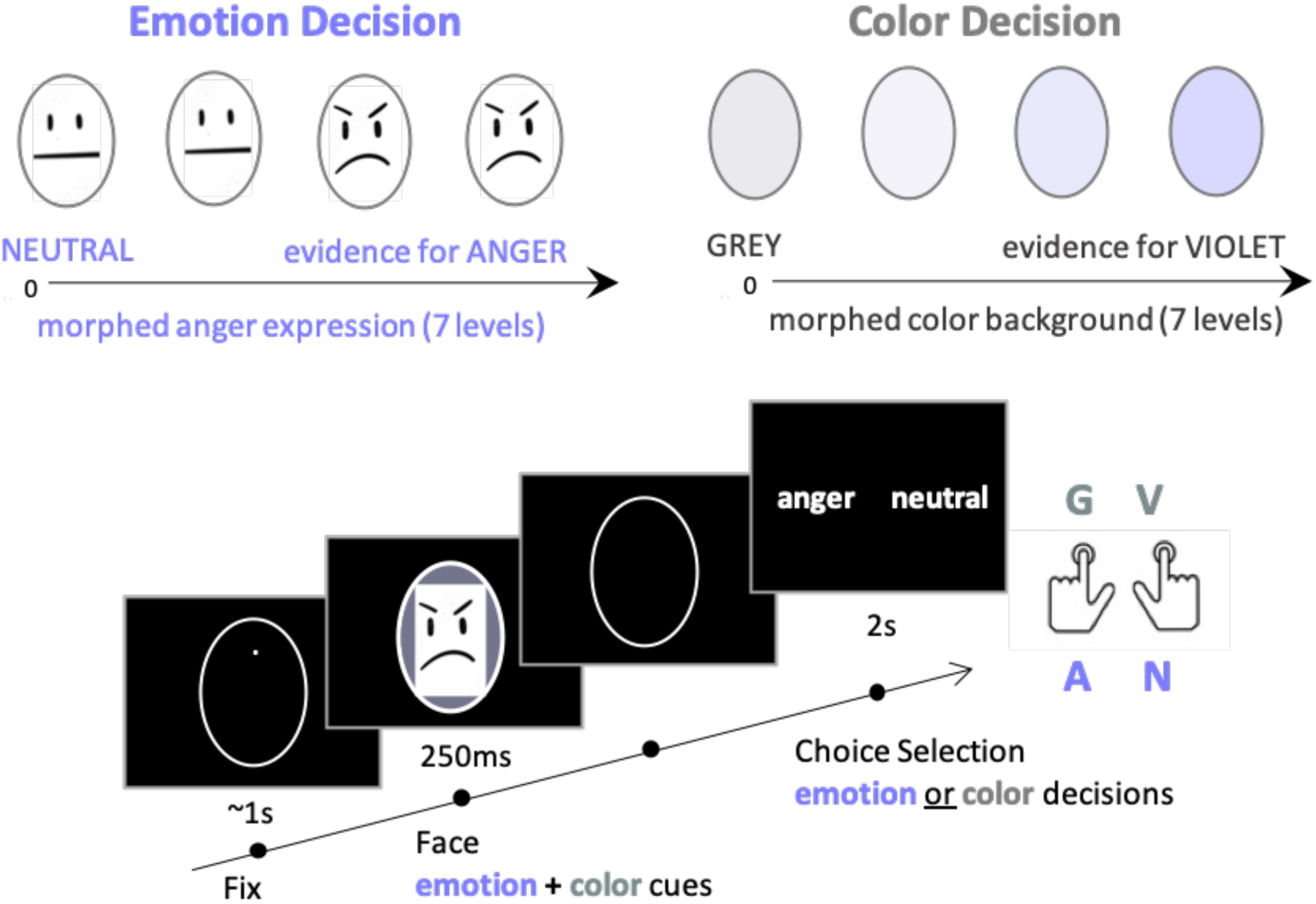
Stimuli and experimental procedure. The top panel shows examples of morphed anger expressions and color masks used to orthogonally manipulate the strength of perceptual evidence in the stimuli. Note that only four of the seven levels of morphs created from neutral to anger and from grey to violet are presented. The bottom panel illustrates the time course of one trial during perceptual detection tasks. After a fixation period, a morphed face, on top of which is superimposed a morphed color mask, was displayed for 250ms. After face offset, and depending on the block, participants reported the presence or absence of either emotion (anger) or color (violet) in the stimuli, while ignoring the other task-irrelevant dimension. For the purpose of BioRxiv submission, natural- appearing face stimuli have been replaced with symbols.

We expected that threat processing would be prioritized to facilitate rapid behavioral adaptation, leading to (Hypothesis 1 – H1) earlier selective encoding of the threat-related dimension in the emotion task, as compared to the encoding of the color dimension in the color task; and (H2) earlier prediction of choice in motor-related signals in the emotion task compared to the color task. Finally, both information processing (H3a) and choice prediction (H3b) should be stronger for high compared to low threat signals, i.e., for angry faces associated with direct gaze compared to averted gaze ^17,36,52,53^_._

## Methods

### Participants

Thirty-eight healthy young adults (20 females; mean age ± s.d. = 23.15 ± 3.38 years) participated in an EEG experiment. All participants were right-handed, had normal vision and no neurological or psychiatric history. The experimental protocol was approved by INSERM and the local research ethics committee (Comité de protection des personnes Ile de France III – Project CO7-28, N° Eudract: 207- A01125-48) and was conducted in accordance with the Declaration of Helsinki. Participants provided written informed consent and were compensated for their participation. One participant was excluded from the analyses due to a technical error while saving the data, and analyses were performed on 37 participants (20 females, mean age = 23.16± 3.42 years).

### Stimuli

The stimuli consisted of morphed facial emotion expressions ranging from neutral to angry expressions ^36,54^ that have been used in several studies ^36,52,55,56^. The full description of the stimuli can be found in El Zein et al. ^36^, and the stimuli are available upon request. 16 identities (8 females) were included in the current study. They varied in gaze direction (direct or averted 45° to the left or right) and emotion intensity – from neutral to angry expressions with 7 levels of emotion. In addition, we created color masks over the faces, morphed from grey to violet with 7 levels of violet. We calibrated the morphing between the emotional expressions and the color masks by performing an intensity rating pre-test of the emotional and color morphs and adjusting the morphs based on the results. Participants (n=10) were presented with the facial expressions or the color masks for 250ms and rated their perceived intensity on a continuous scale from “not at all intense” to “very intense” using a mouse device (with a maximum of 3 seconds to respond). We adjusted for differences between emotion and color morphs by linearizing the mean curves of the judged intensities and creating corresponding morphs that were equalized according to the perceived intensities.

### Experimental paradigm

Using the Psychophysics-3 toolbox ^57,58^, stimuli were projected onto a black screen. Each trial began with a white oval delineating the faces that remained throughout the trial (see **Figure 1**). The white oval appeared for approximately 500ms, followed by a white fixation dot presented at the eye level for approximately 1000ms (to ensure a natural fixation on the upcoming faces and to avoid eye movements from the center of the oval to the eye region). Then, the morphed face, over which we superimposed a morphed color mask, was presented for 250ms. After the face offset, participants had to make either an emotion or a color decision in a blocked design. Depending on the block, participants were asked to report (max. 2s after face offset) the presence or absence of either emotion (anger) or color (violet) in the stimulus, while ignoring the other task-irrelevant dimension (see Figure 1). They provided their response by pressing one of the two buttons ‘Right Control’ and ‘Left Control’ keys on the keyboard with their right and left index, respectively. A grey/violet and neutral/anger key mapping was used (4 possible hand mappings), kept constant within subjects and counterbalanced across subjects. The experiment was divided into 32 experimental blocks (16 blocks of emotion decisions, 16 blocks of color decisions), each consisting of 32 trials, balanced in the number of gaze directions (2), gender (2), and morph levels (8). Color masks and emotions were combined semi-randomly, such that within each block, no correlation between emotion and color intensity was allowed (r<0.05, p>0.6). This resulted in a total of 512 trials (32 trials*16 identities) per decision type (emotion/color), and 1024 trials for the entire experiment. Participants alternated between the emotion and color tasks at each block. Prior to the experiment, each participant completed a short practice session (one emotion and one color block, two identities, 16 trials each).

To ensure that the observed differences at the neural level between tasks did not depend on differences in difficulty, we equalized detection sensitivity across emotion and color dimensions using an adaptive Bayesian titration procedure. Participants completed a short titration session of two blocks (one emotion and one color block, two identities, 32 trials each), and each participant’s perceptual sensitivity to both emotion and color dimensions was estimated separately. The choices of each participant were regressed against the emotion or color morph level in a general linear model (GLM), and parameter estimates were extracted for each dimension. These parameter estimates were then used to calculate a titration factor (i.e., multiplier) that was multiplied by the color mask morph applied to the face, allowing the color level of the stimulus to be rescaled to the same perceptual intensity as the emotion level ^59^. This procedure was reiterated every two blocks throughout the experiment. This ensured that the difficulty between emotion and color decisions was calibrated and updated across the experiment, so that the same slope of the psychometric function reflected participants’ behavior in both conditions.

Participants were informed prior to the experiment that they would receive one point for each “correct” response (i.e., correct detection when the morph level was >3), which could result in a bonus of 5 to 10 Euros, depending on their final score. Feedback on their performance (i.e., percentage of correct responses) was displayed on the screen after each block. Every 8 blocks, a progress bar allowed participants to estimate how close they were to receiving a bonus. At the end of the experiment, all participants received the same compensation, regardless of their performance.

### Behavioral data analyses

The theoretical framework of Signal Detection Theory distinguishes between sensitivity to sensory information and response bias (or criterion) that reflects the observer’s tendency to interpret the face as displaying either one of the two options (anger or neutral for emotion decisions, violet or grey for color decisions). Within this framework, we used a model of choice hypothesizing that decisions are formed based on a noisy comparison between the displayed emotion or color and a criterion. For emotion and color decisions separately, we fitted the data with the simplest model (model 0) that could account for each subject’s decisions using a noisy, ‘signal detection’-like psychometric model, to which we included a lapse rate to account for random guessing:

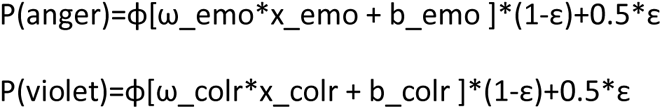

where P(anger) and P(violet) correspond to the probability of judging the face as angry or violet, respectively, ɸ(.) to the cumulative normal function, ω_emo and ω_colr to the perceptual sensitivity to the displayed emotion or color, x_emo and x_colr to the a trial-wise array of evidence values in favor of anger (emotion intensity in the stimulus) or of violet (color intensity in the stimulus) (from 0 for neutral to +7 for an intense display), b_emo and b_colr to an additive, stimulus-independent bias toward one of the neutral/anger or grey/violet choices, and ε to the proportion of random guesses among the choices. Sensitivity (ω_emo, ω_colr) and bias (b_emo, b_colr) parameters computed with these models were extracted and compared using t-test analyses. We also assessed “interference" effects between stimulus dimensions by examining whether the decision-irrelevant stimulus dimension (i.e., emotion intensity in color decisions, and color intensity in emotion decisions) influenced the processing of decision-relevant cues.

### Drift-diffusion model analyses of the behavioral data

To better understand the difference in decision bias parameters and reaction times between the Emotion and the Color tasks (see behavioral results), we fitted DDMs choices and RTs distributions using the Python package HDDM ^60^. HDDM implements a Bayesian hierarchical estimation of DDM parameters at the individual and group level using Markov Chain Monte Carlo (MCMC) sampling. DDMs describe decision processes in dichotomous choice tasks by several parameters representing different aspects of the decision, including: a drift rate (v), which describes the rate at which evidence for one of the choices accumulates; a threshold (a), which represents the amount of information required to make a choice (i.e. the distance between the two response boundaries); a pre-decisional bias (or starting point bias; z), representing the starting point of the decision between the two alternatives, closer (more biased, requiring less evidence to choose) or further (less biased, requiring more evidence) from one alternative; a non-decision time (t0), representing the portion of the response time not involved in a decision process but in stimulus encoding and response execution. To fit the models, responses were coded “Angry/Violet” responses and “Neutral/Grey” responses during the Emotion and Color tasks, respectively. The task (Emotion vs Color), the continuous intensity level (centered, –3.5 to 3.5 in eight steps), and their interaction were fitted as regression parameters in the DDMs. To improve the interpretability of the parameter estimates, we adopted a cell design matrix estimating one parameter (a, v, z or t0) for each task, and the effect of intensity on each of them.

Eight models were run to understand the influence of task type and intensity level on decision. First, we ran 7 models to test the effect of task and intensity on drift rate (v), threshold (a), and non-decision time (t0) individually or in combination. In all these models, we estimated separate bias (z) parameters by task. Finally, we tested whether the best-fitting model in the first step of model selection was improved by removing the effect of task on the bias parameter. Due to the moderate number of trials per condition, we simplified the models in several ways: the boundaries of the model were associated with “Angry/Violet” (upper threshold) and “Neutral/Grey” responses (lower threshold), regardless of whether they required a left or right button press (correspondence was counterbalanced across participants, but stable within participants); furthermore, the inter-trial variabilities were fixed to zero, since a proper fit of these parameters is particularly challenging, especially with small to medium-sized trials number ^61^.

Models were compared using their Deviance Information Criterion (DIC) and Bayesian Predictive Information Criterion (BPIC) values ^62^. Further analyses were performed on the best-fitting model (i.e., lower DIC and BPIC). We based inference on the 95% credible intervals (95%CrI) of the group-level parameters, defined as the interval between the 2.5^th^ and 97.5^th^quantiles of the estimates’ posterior distributions (or the posterior distribution of the parameter differences when testing for them). We considered a parameter (or a difference) to be credible if the 95%CrI did not overlap with 0 (or 0.5 in the case of the starting point bias, i.e., a perfectly equidistant point between the two response boundaries). MCMC sampling was generated using 3 chains of 5000 iterations each (1500 of which were used as a burn-in period). Detailed information about model diagnostic can be found in Supplementary Text.

### EEG acquisition and pre-processing

EEG was recorded continuously from 64 scalp sites with CMS/DRL reference electrodes using a BioSemi headcap with active electrodes. The EEG signal was amplified using an ActiveTwo AD-box amplifier (BioSemi), low-pass filtered online (250 Hz) and digitized at 1000 Hz. Raw EEG data were pre- processed using the Brainstorm toolbox for MATLAB ^63^.

First, raw EEG was recalculated to average reference, down-sampled to 500 Hz (2ms steps), and band-pass filtered between 1 and 40 Hz. Second, EEG data were visually inspected to remove muscle artifacts and to identify noisy electrodes, which were interpolated to the average of adjacent electrodes. Third, independent component analysis (ICA) excluding interpolated electrodes was performed on the continuous data and ICA components capturing eye blink artifacts were rejected. These EEG data were then epoched from 2s before to 2s after the face stimulus onset (20ms steps) and linearly de-trended. Epochs containing activity exceeding a threshold of +/- 70 µV were automatically discarded. Finally, the resulting individual epochs were visually inspected to manually exclude any remaining trials with artifacts. After trial rejection, the remaining trials averaged 975 ± 46 trials per subject. The resulting data were resampled to 100 Hz (10ms steps) from 500ms before to 1.5s after stimulus onset.

Time-frequency decomposition was also performed using a Brainstorm Toolbox pipeline. Time-frequency representations (TFRs) of individual trials were computed for each subject. The Morlet Wavelet transformation with 3-s time resolution at the central frequency 1 Hz (as calculated by the full width at half maximum; FWHM) was used to calculate spectral power estimates at each point of the time-frequency window ranging from –2s to 2s (20ms steps) in the time domain and 1 to 40 Hz (1 Hz steps) in the frequency domain. TFRs were baseline corrected with respect to the power averaged over the entire epoch.

### EEG analyses

#### Neural encoding of emotion and color intensity

To test Hypothesis 1 of an earlier selective encoding of threat-related information during the emotion vs the color task, we applied single-trial EEG regressions (general linear models) against the intensity of the displayed emotion or color mask at each electrode and time point following stimulus presentation ^35,64^. We added the choice as an additional regressor to control for the participants’ subsequent detection reports (detection of violet in the color task, detection of anger in the emotion task) on a trial-by-trial basis. The resulting time course at each electrode represents the degree to which the EEG activity ‘encodes’ (co-varies with) the emotion or color strength provided by the morphed facial features or color intensity. We conducted this analysis separately for the emotion task (regressor: emotion intensity) and the color task (regressor: color intensity) and performed a multiple comparison correction on the parameter estimates against zero to isolate the electrodes and timepoints where encoding of intensity was significant (**Table 1**). To do so, we used Fieldtrip’s multiple comparison analysis ^65^, using the MonteCarlo method to calculate the probability of significance, dependent sample T-statistics, cluster corrections (clusteralpha = 0.005, cluster statistic =maxsum, correct tail=prob, number of permutations=1500). We also conducted this multiple comparison analysis on the difference between the parameter estimates for emotion encoding and color encoding (see **Table 1**). Finally, to test our Hypothesis 3a, we did the same analysis for emotion encoding in the emotion task, this time isolating facial expressions with a direct gaze and those with an averted gaze (see **Table 2**). As in the emotion vs. color tasks, here we corrected for multiple comparisons for the effects against zero, and for the differences between the two conditions (gaze direct vs gaze averted).

**Table 1.**
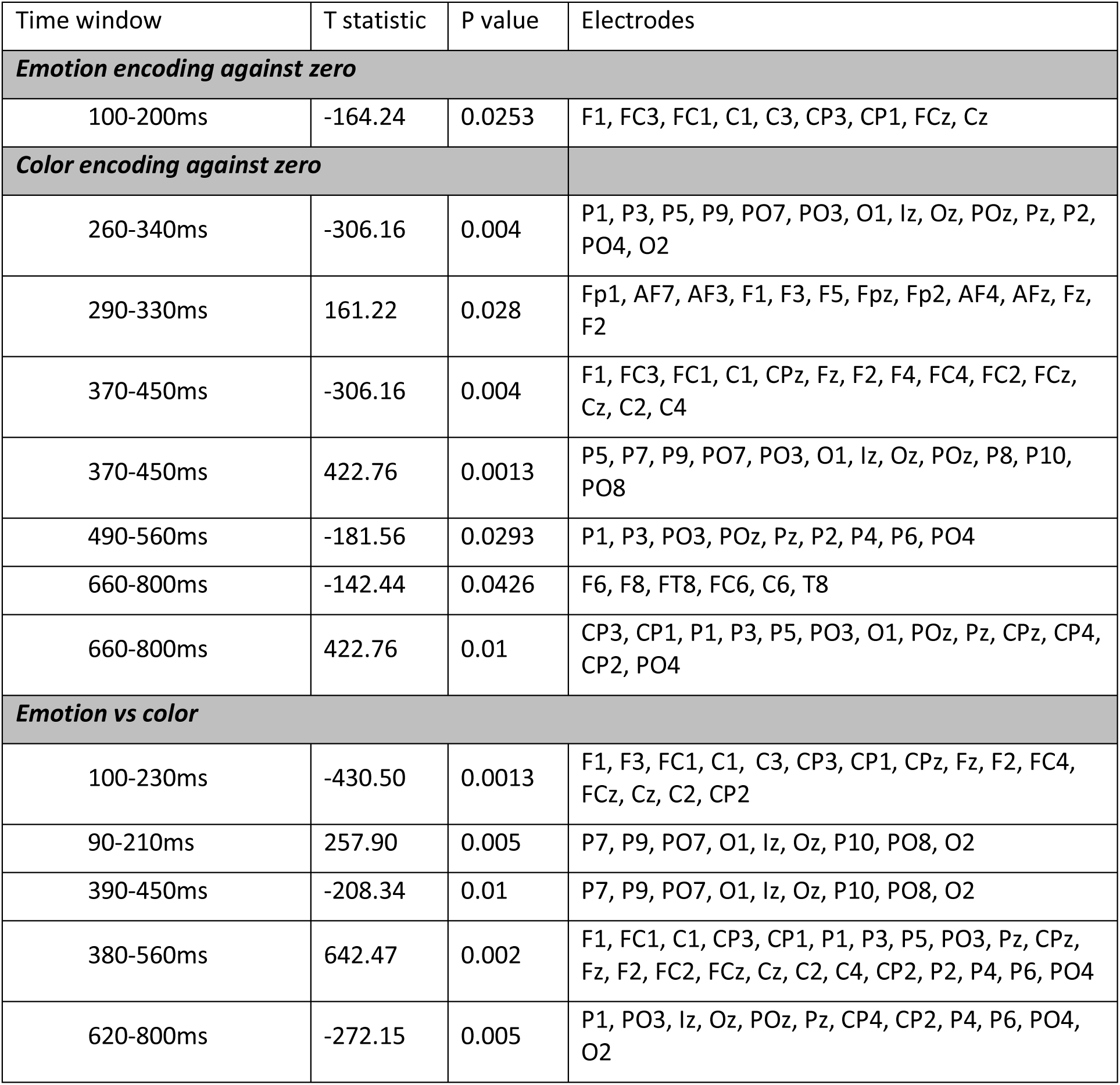
Significant clusters for the encoding of emotion and color evidence.

**Table 2.** Significant clusters for the encoding of emotional expressions with direct and averted gaze.

| Time window | T statistic | P value | Electrodes |
| --- | --- | --- | --- |
| ***Direct gaze emotion encoding against zero*** |  |  |  |
| 80-130ms | -114.65 | 0.0706 | F1, FC3, FC1, C1, C3, Fz, F2, FCz |
| 270-330ms | 152.96 | 0.03 | AF3, F1, F3, Fpz, AF4, AFz, Fz, F2 |
| 270-320ms | -129.05 | 0.056 | P5, P9, PO7, PO3, O1, Oz, P8, P10, PO8, PO4, O2 |
| ***Averted gaze emotion encoding against zero*** |  |  |  |
| None |  | p>0.9 |  |
| ***Direct vs averted gaze emotion encoding*** |  |  |  |
| None |  | p>0.6 |  |

#### Prediction of choice with motor lateralization

To test Hypothesis 2 of an earlier prediction of choice in motor-related signals during the emotion vs. the color task, we assessed whether and when the detection of choice can be predicted by the motor lateralization index in the mu-alpha (8-12 Hz) and –beta (12-32 Hz) frequency bands (^30^.

To compute the motor lateralization index, we first isolated effector-selective (left vs right hand) neural activity. As mu-beta activity is known to be suppressed at response time in the hemisphere contralateral to the hand used for response ^30,66^, we calculated the spectral power from 8 to 12 Hz or 12 to 32Hz at response time for each electrode and time point for all subjects, for both emotion and color decisions. The resulting response-locked mu-alpha or beta activity for the trials in which subjects responded with their right hand was then subtracted from that of trials in which subjects responded with their left hand. After averaging across all subjects, the electrodes where the motor lateralization was maximal at 200ms before response time were identified at ’P3,’CP3’,’C3’ for the left hemisphere and ’P4,’CP4’,’C4’ for the right hemisphere for both types of decisions. For emotion/color decisions, motor lateralization specific to ‘anger’/’violet’ responses was obtained by subtracting contralateral from ipsilateral mu spectral activity (8-12Hz or 12-32Hz) relative to the hand assigned to ‘anger’/’violet’ responses (counterbalanced across subjects), over effector-selective electrodes at each time point for all subjects. We then performed a logistic regression to predict choice (Anger/Neutral in the emotion task, Violet/Grey in the color task) with the motor lateralization index as predictor, and trial-by-trial reaction times as an additional regressor to isolate for effects that are independent of participants’ fluctuations in response times and differences in RTs between the two tasks.

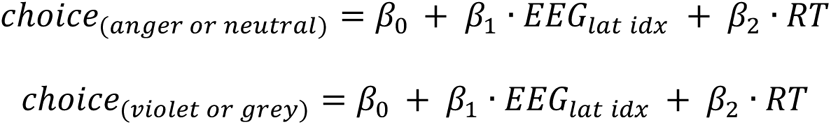

We extracted and plotted the parameter estimates of this regression over time (see **Figure 3c**) and compared the parameter estimates predicting the response Anger with the parameter estimate predicting the response Violet at each time point using a paired samples t-test. We corrected for multiple comparisons by randomly shuffling the pairings between responses and EEG signals 1000 times. The maximal cluster-level statistics (the sum of t-values across contiguous significant time points at a threshold level of 0.05) were extracted for each shuffle to compute a ‘null’ distribution of effect size across the [-200, +1100]ms time window. For the significant cluster in the original data, we computed the proportion of clusters in the null distribution whose statistic exceeded that obtained for the cluster in question, corresponding to its ‘cluster-corrected’ p.

**Figure 3.**
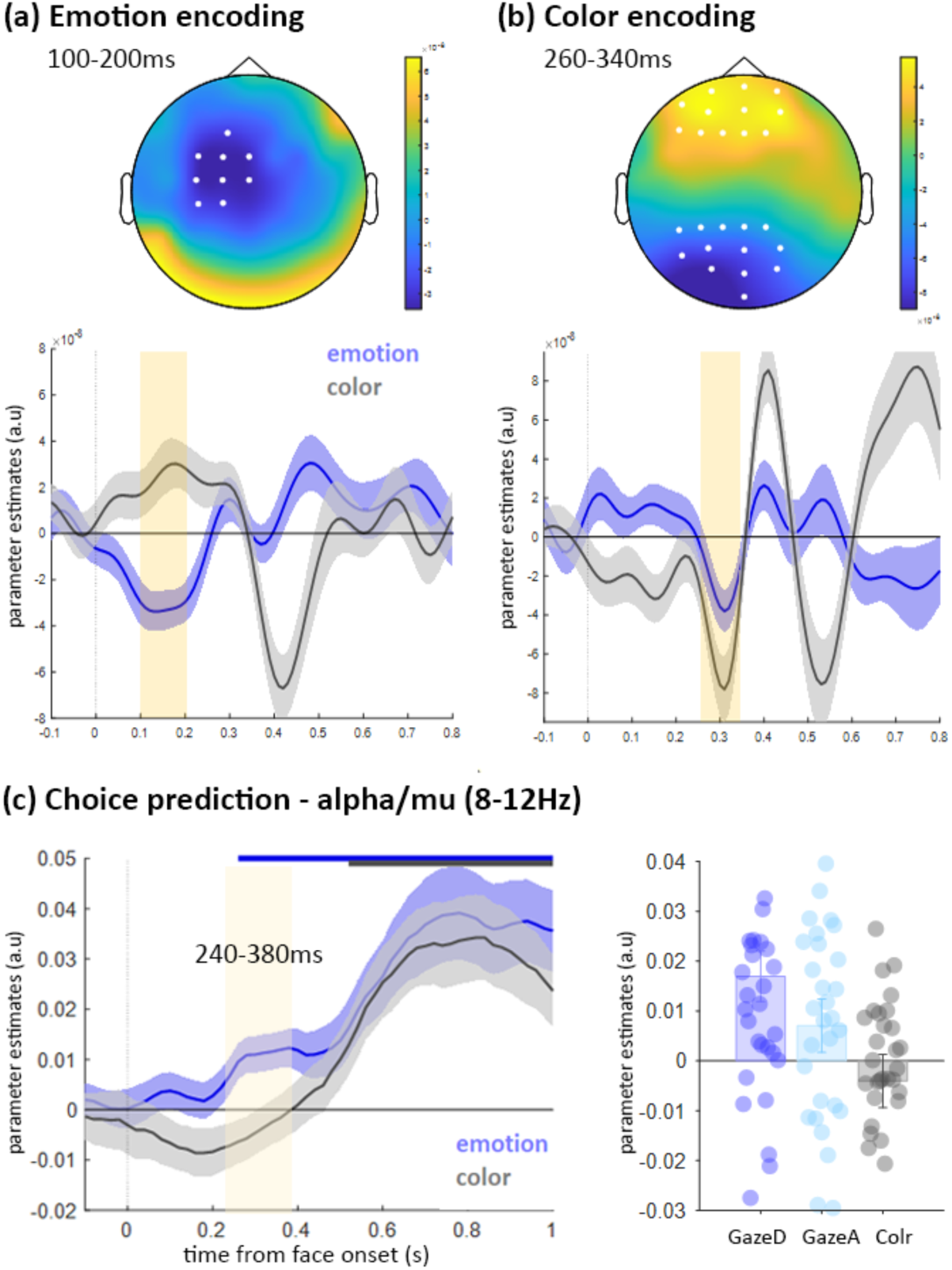
Neural encoding and choice-predictive activity. **(a)** Scalp topography of the first significant cluster for the encoding of emotion intensity (against zero), that emerged as soon as 100-200ms on frontal and central electrodes. Significant electrodes are highlighted in white. A similar significant cluster was found when emotion encoding was contrasted to color encoding (see Table 1). Below the scalp topography, encoding time course for emotion (blue) and color (grey) decisions separately, expressed as mean parameter estimates in arbitrary units (a.u.) at electrodes of interest. Shaded error bars indicate s.e.m. Shaded light yellow area indicates the significant cluster time window (100- 200ms). **(b)** Scalp topography of the first significant cluster for the encoding of color intensity (against zero), that appeared at 260ms on parietal and frontal electrodes. Same conventions as (a). **(c)** Time course of choice-predictive activity (stimulus-locked) in the motor lateralization index for emotion and color decisions in mu frequency band (8-12Hz). Note that the motor-preparatory EEG signals were predictive of participant choices from 240ms after stimulus onset during emotion decisions, and only from 520ms following stimulus presentation during color decisions. Full lines are the times where the parameter estimate is significant against zero (blue emotion task, grey color task). Shaded light yellow area indicates the timepoints where there is a significant difference between the two (240ms to 380ms). The left panel shows the extracted parameter estimates averaged across the motor lateralization electrodes over the 240-380ms window, separately for faces with direct (GazeD) and averted (GazeA) gaze.

To test our hypothesis 3b, we predicted the response ‘anger’ with the motor preparation measure in the mu-alpha band on the mean activity of the isolated significant cluster from 240ms to 380ms, separately for direct and averted gaze.

## Results

### Behavioral results from the emotion vs. color task

Our first hypothesis predicted that emotion intensity encoding in the emotion task would be observed earlier than color intensity encoding in the color task over both sensory and motor brain areas. To ensure that this difference does not stem from different difficulties in the two tasks, we used a Bayesian titration procedure equalizing detection sensitivity across both tasks. This procedure was successful as no differences in sensitivity were observed between the two tasks (see Methods for the choice model used to compute sensitivity): sensitivity for the emotion task (0.63±0.16), sensitivity for the color task (0.64±0.17), no difference between the two (T_37_=0.33, p=0.73, BF=0.187) (**Figure 2a** – left panel). However, there was a difference in the bias parameter between the emotion task (- 2.69±0.64) and the color task (-3.16±0.99), difference between the two (T_37_=2.40, p=0.02), and we therefore accounted for the choice at each trial in our encoding regressions. Finally, participants were faster in the color task (mean ± sem: 690±39ms) as compared to the emotion task (mean ± sem: 820±46ms) (T_36_=-9.280, p<0.001) (**Figure 2a** – left panel).

**Figure 2.**
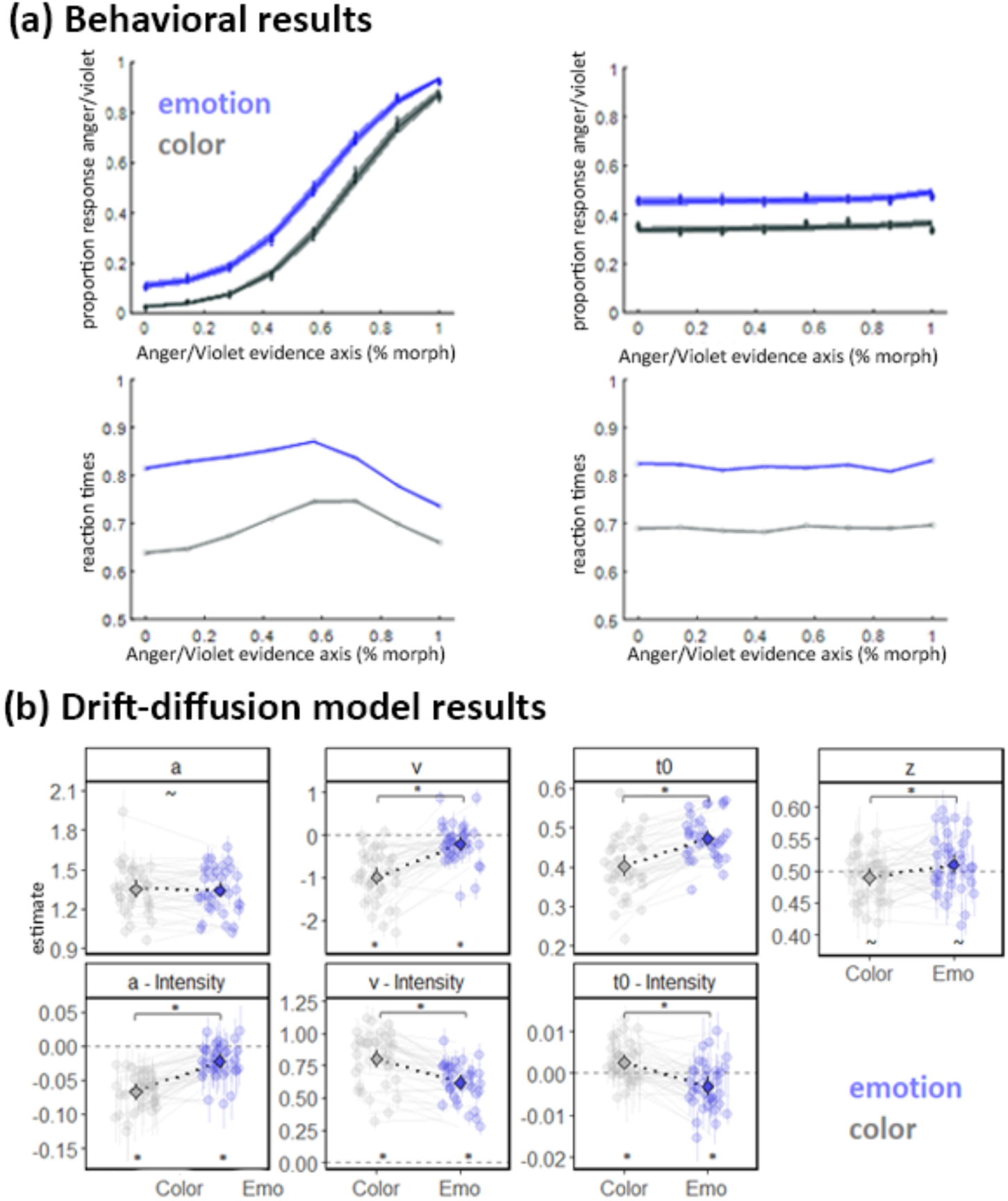
Behavioral results. **(a)** The top-right panel shows the psychometric function representing the proportion of ‘anger’ (in blue) or ‘violet’ (in grey) responses as a function of the strength of the decision-relevant perceptual evidence for anger or violet (percentage of morph, 0 = neutral or grey, 100% anger or violet). Dots and attached error bars indicate the experimental data (mean ± sem). Lines and shaded error bars indicate the prediction of the fitting model. The bottom-right panel represents mean reaction times as a function of the strength of perceptual evidence for anger or violet. Top– and bottom-left panels present the Psychometric function representing the proportion of ‘anger’ or ‘violet’ responses (top) and mean reaction times (bottom) as a function of the strength of the decision-irrelevant perceptual evidence (i.e., emotion intensity in color decisions, and color intensity in emotion decisions). **(b)** Drift-diffusion model results. The best-fitting model included the effect of task, intensity level, and their interaction on threshold (a), drift rate (v), and non-decision time (t0), and an effect of task on starting point bias (z). Light points and lines: Individual estimates and 95% Credible Interval. Dark points and lines: Group level estimates and 95% Credible Interval.

To test for interference effects, we also examined whether responses and reaction times changed as a function of the irrelevant dimension (color intensity in the emotion task, emotion intensity in the color task). There was no interference, as the percentage of anger choices did not vary with color intensity nor did the associated reaction times, and the percentage of violet choices did not vary with anger intensity, nor did the associated reaction times (**Figure 2a** – right panel).

### Drift-diffusion model results

To better understand the difference in reaction times between the Emotion and the Color tasks, we compared different drift-diffusion models. The best-fitting model included the effect of task, intensity level, and their interaction on threshold (a), drift rate (v), and non-decision time (t0), and an effect of task on starting point bias (z) (see **Figure 2b**). The three chains converged properly (maximum R-hat = 1.009; minimum Effective Sample Size = 1026; the ‘hairy caterpillar’ shape was visually verified), and the simulated data showed a high correlation with the observed data (Anger/Violet response proportion: rcol = .998, remo = .998; Anger/Violet RTs: rcol = .955; remo = .977; Neutral/Grey RTs: rcol = .989; remo = .985)(see Supplementary Text for detailed information about model diagnostic).

Posterior parameters testing on the starting point bias (z) indicated a credible small difference between the Color and Emotion tasks (zcol-emo = –.021; 95%CrI = [-.041, –.0001]; p>0 = .024), with a lower starting point in the Color task compared to the Emotion task. However, neither the starting point for the Color task (zcol = .489; 95%CrI = [.475, .503]; p>.5 = .058) nor that for the Emotion task (zemo = .510; 95%CrI = [.494, .525]; p>.5 = .896) were credibly different from 0.5. This pattern suggests that even if participants were slightly more likely to choose Angry rather than Neutral compared to Violet rather than Grey, they did not exhibit a credible response bias in either task.

Posterior parameters testing on the threshold parameter (a) showed no credible difference between tasks (acol-emo = .008; 95%CrI = [-.079, .098]; p>0 = .574). However, the continuous effect of Intensity was credible in both the Color task (βcol = –.068; 95%CrI = [-.080, –.056]; p>0 = 0) and the Emotion task (βemo = –.023; 95%CrI = [-.034, –.012]; p>0 = 0), with greater intensity being associated with a lower threshold in both tasks. Furthermore, the effect of intensity was credibly more negative in the Color task compared to the Emotion task (βcol-emo = –.045; 95%CrI = [-.061, –.028]; p>0 = 0), suggesting that the lowering of response threshold with increasing intensity level was present in both tasks, but with a stronger effect in the Color task.

Posterior parameter testing on the drift rate (v) revealed that both the drift rate for the Color task (vcol = –.987; 95%CrI = [-1.194, –.778]; p>0 = 0) and for the Emotion task (vemo = –.211; 95%CrI = [-.370, –.048]; p>0 = .006) were credibly lower than 0. Furthermore, the drift rate in the Color task was credibly lower than in the Emotion task (vcol-emo = –.778; 95%CrI = [-1.038, –.510]; p>0 = 0). Given that these estimates represent the parameter at the level 0 of intensity, these results suggest that for these ambiguous stimuli, participants favored the decision toward the Grey and Neutral boundaries. However, we additionally found a credible effect of Intensity for both the Color (βcol = .804; 95%CrI = [.731, .876]; p>0 = 1) and the Emotion task (βemo = .619; 95%CrI = [.565, .672]; p>0 = 1), and a credible difference of these effects between the tasks (βcol-emo = .186; 95%CrI = [.094, .275]; p>0 = 1), with a stronger effect of intensity in the Color task. Therefore, as expected, increasing morph intensity toward a Violet or Angry stimulus led participants to more efficient evidence sampling toward that boundary and a greater evidence integration in the Color task.

Finally, posterior parameter testing on the non-decision time (t0) showed a credible difference in the non-decision time between the tasks (t0Col-Emo = –.070; 95%CrI = [-.103, –.037]; p>0 = 0), with more time required to encode the stimulus and/or execute the response in the Emotion task. Interestingly, we found the Intensity effect to be credible for both the Color and for the Emotion tasks. However, while non-decision time increased with intensity for the Color task the (βcol = .003; 95%CrI = [.001, .004]; p>0 = .994), it decreases with intensity for the Emotion task (βcol = –.003; 95%CrI = [-.005, –.001]; p>0 = .005), as further confirmed by a credible posterior difference between these two effects (βcol = .006; 95%CrI = [.003, .009]; p>0 = 1).

### Neural encoding of emotion and color intensity

We assessed whether encoding of emotion intensity in the emotion task emerged earlier than the encoding of color intensity in the color task, over both sensory and motor brain areas (Hypothesis 1). In the emotion task, one significant cluster for encoding emotion intensity emerged as soon as 100- 200ms on frontal and central electrodes (see **Table 1** and **Figure 3a**). In the color task, seven significant clusters appeared. The earliest appeared at 260ms on parietal and frontal electrodes (see **Figure 3b**). The other five clusters emerged at 370ms, 490ms and 660ms (see **Table 1** for full details). When looking for significant clusters of the difference in encoding between emotion and color intensity, the early cluster corresponding to emotion encoding did indeed appear at 100 to 230ms. For the color encoding, the earliest cluster at 290ms did not appear when directly comparing emotion and color encoding, but those at 370ms and 660ms did. In conclusion, there is a specific early encoding of emotion intensity around 100ms, versus a later encoding of color information starting at 270ms.

### Choice prediction in the emotion vs color task

Our second hypothesis predicted that the perceptual choice would be predicted earlier in the motor-related signals during the Emotion task compared to the Color task ^17,36^. We performed a

logistic regression to predict the choice of Anger or Violet in the Emotion and Color tasks, respectively, with motor signals in the 8-12Hz mu-alpha band, and reaction times as an additional regressor. The anger response was significantly (t-test against zero, p<0.05) predicted as soon as 260ms, while the violet response was predicted at 520ms, both up to 1sec after stimulus onset (difference between the 2 significant from 240ms to 380ms at p<0.05, cluster-corrected p-value<0.001, **Figure 3c**).

We performed the same analysis with the motor signals in the 12-32Hz mu-beta band. Here, there were no differences in choice prediction between the two tasks (all p>0.08). The prediction of choice by the motor signals was significant for both the anger response and the violet response around 420ms after stimulus onset (starting at 420ms for the emotion task, and at 440ms for the color task, p<0.05

### High threat signals influence on encoding and choice prediction

Our third hypothesis predicted that both the encoding (H3a, Hypothesis 1 with the gaze direction) and the choice prediction (H3b, Hypothesis 2 with the gaze direction) should be stronger for high vs. low threat signals, i.e., for angry faces associated with direct gaze vs. averted gaze. To test whether gaze direction influenced emotion processing (H3a), we ran a cluster analysis on the encoding of emotion associated with direct and averted gaze separately, and then a direct comparison of the two. Only close to significance effects emerged for direct gaze emotion encoding (see **Table 2**), suggesting that the effects may be more important for high threat signals (although no significant differences between direct and averted gaze were observed).

To test whether choice prediction was stronger for angry faces with a direct gaze, we predicted the response ‘anger’ with motor preparation measure in the mu-alpha band in the previously isolated time window from 240ms to 380ms, separately for direct and averted gaze. We found that only the choice prediction for faces with a direct gaze was significant against zero (T_36_=3.52, p=0.001), while it was not significant for faces with an averted gaze and for the color choice (p>0.17). There was a significant difference between the emotion choice prediction when the faces displayed a direct gaze as compared to the color choice prediction (T_36_=3.37, p=0.0018), but there were no significant differences between choice prediction for gaze direct vs. averted gaze, nor for averted gaze vs. color (p>0.10) (**Figure 3c**).

## Discussion

The present study investigates whether perceptual decisions about the emotional, threat-related aspects of stimuli engage specific or similar neural computations compared to decisions on their neutral, non-threatening components. We used electroencephalography (EEG) to simultaneously record brain activity, while participants performed two different detection tasks (emotion or color) on the same, two-dimensional visual stimuli. Detection sensitivity was equalized across dimensions using an adaptive titration procedure. Our results revealed three specific effects when perceptual decisions concern emotion rather than color. First, the amount of perceptual evidence (intensity) was encoded earlier (100ms) in a cluster of central electrodes in the emotion task compared to the color task (290-330ms). Second, participants’ choices were predicted earlier by the mu-alpha EEG rhythm in the emotion task (240ms) than in the color task (380ms). Third, this choice-predictive activity over motor cortex was stronger for high-threat signals, i.e., for angry faces with direct versus averted gaze. Taken together, these findings indicate that perceptual decisions regarding the threat-signaling dimension of facial displays are associated with prioritized neural coding in action-related brain regions, supporting the motivational value of socially relevant signals.

### Encoding of emotion and color intensity

Consistent with previous findings that threatening facial expressions receive privileged processing compared to neutral expressions and non-social stimuli ^e.g. 6^, we observed earlier (100-200ms) encoding of evidence strength in the emotion task compared to the color task (290-330ms). Past experiments have shown that electrophysiological (MEG or EEG) activity discriminates emotional (anger) expressions from neutral expressions in the first 100ms ^67^ and correlates with BOLD responses in the amygdala and occipital cortices ^68^. Moreover, in contrast to gender or identity tasks, which are restricted to occipital regions, early emotion decoding (110-210ms) involved a widely distributed network, including both temporo-parietal and frontal regions ^69^. Together, these studies show that emotional, and particularly threatening, facial expressions are processed rapidly (from around 100ms) by both subcortical and cortical brain networks. Here, by contrasting two perceptual decision tasks (emotion and color) performed on the same two-dimensional visual stimuli, we show that privileged processing in such an early time window (100-200ms) benefits only the emotional, threat-signaling dimension.

Several authors have argued that perceptual saliency (including low-level perceptual features) and appraisal relevance jointly drive prioritized stimulus processing ^70–73^. In the present study, owing to a titration procedure that equalized detection sensitivity (and thus presumably task difficulty) across dimensions, highly salient information in the stimuli (angry expressions) did not interfere with the processing of color information during color decisions (see **Figure 2a**). Prioritization of the emotional, threat-related dimension may thus only occur when this information was relevant to the task at hand (i.e., during emotion decisions, not color decisions, see also ^74^), concurring the suggestion that relevance detection is not automatic, but depends on task demands such as perceptual and/or cognitive load ^72^. Besides being task-relevant, the emotional, threat-related dimension also varies in terms of its immediate relevance to the observer. Indeed, angry faces associated with direct or averted gaze clearly differ in terms of implied threat to the observer and have been shown to be appraised as high or low threat, respectively ^e.g. 48–50^. Surprisingly, and contrary to our hypothesis based on previous findings ^36^, no significant difference was observed between direct and averted angry faces in their early perceptual processing in the present study. Only near-significant effects were found for direct gaze emotion encoding. Note that this absence of significant difference between direct and averted angry displays may be related to a lack of statistical power, as only half of the trials were presented as compared to El Zein et al. ^36^.

#### Choice prediction

Additionally, and similar to past experiments that observed a buildup of choice-selective activity over motor-related regions during perceptual decision on non-social stimuli ^30,31,35,66,75–78^, it was possible to statistically predict participants’ choices from the mu/alpha power over motor cortical areas. This signal has been suggested to reflect the formation of an upcoming decision, i.e., a commitment to one choice alternative ^30,32,79^. Here, we showed that the onset of choice predictive activity was earlier for emotion decisions (260ms after stimulus onset) than for color decisions (500ms), strongly suggesting that the emotional, threat-related dimension of our complex two-dimensional stimuli was not only processed, but also translated into action selection earlier than the color dimension. Moreover, this earlier choice predictive activity was stronger for high-threat signals, i.e., for angry faces with direct gaze compared to averted gaze.

The question remains as to why perceived social threats, notably those that are observer- relevant, lead to earlier selection between action alternatives in motor-related cortices. One likely explanation is related to the high behavioral/motivational relevance of emotional expressions. Evolutionary accounts of emotional displays argue that the very function of emotions is to serve communicative purposes by conveying critical information about the emitters ^80–82^ while also facilitating behavioral responses in the observers ^83,84^. Emotional faces can be translated into action possibilities via different neural pathways ^see^ ^review^ ^85^. In humans, a growing body of literature concurs with evolutionary accounts by highlighting a functional and anatomical link between neural systems that sustain emotional appraisal and those that underlie action preparation ^21,86–88^. Moreover, and in agreement with the present results, the influence of threat displays on motor-related areas was observed from 150ms to 300ms after stimulus onset ^e.g. 16,17,20,36,52^. We therefore propose that the earlier choice predictive activity during emotion decisions is related to the behavioral relevance of the perceived threat to the observer, rather than to its sensory properties. Threat-signaling emotions would facilitate the selection of adaptive behavioral responses.

Nevertheless, and somewhat counterintuitively, although participants’ choices could be predicted earlier in the emotion task, their reaction times were longer than in the color task (even though we controlled for RTs in the choice predictive analysis). In the past, longer reaction times to emotional stimuli have been suggested to reflect an interference between the attentional resources devoted to their privileged processing and task demands ^89,90^. Slower RTs to threatening stimuli could also reveal momentary freezing, which facilitates risk assessment by enhancing perceptual and attentional processes to the source of danger and preparing subsequent actions ^91,92^. Here, we used drift-diffusion model analysis to show that the emotion task was associated with a smaller decrease in boundary separation with increasing intensity (a) and longer non-decision times (t0) compared to the color task. Consistent with our findings, the presentation of threat-related (fearful) facial expressions led participants to subsequently respond cautiously, as indexed by both slower reaction times and higher boundary separation, compared to neural expressions ^93^. Moreover, higher boundary separation has also been found for emotional faces that have an immediate relevance to the observer ^94^, i.e., fearful faces associated with averted gaze, which are rated as being both more intense ^e.g. 49,50^ and more negative ^95^ compared to direct fearful faces. Beaurenaut et al. ^94^ proposed that participants are particularly cautious about misinterpreting such danger-related social signals. Furthermore, it has been shown that during perceptual decisions on non-social neutral stimuli, both higher boundary separation and larger non-decision time estimates are observed when participants are instructed to trade speed for accuracy ^96^. Thus, in order to be more accurate, participants can increase their response caution, and/or adopt strategic motor slowing, i.e., delay their motor response once their decision has been made ^97^. Even after the decision has been reached, longer t0 in response to threatening signals may suggest that a freezing mechanism is in place, involving motor execution speed. This interpretation seems consistent with the notion of freezing as a state of motor immobility, accompanied by heightened vigilance and caution ^98^. Overall, this suggests that while perceptual decisions about threat-signaling emotions (relative to color) facilitate action selection among available alternatives, participants remain cautious and freeze before fully committing to and implementing the selected plan, to avoid costly misinterpretations.

#### Limitations and conclusion

The present experiment is not without limitations. First, our main results depend on the contrast with a control condition (color) that is neither social nor threatening, and thus cannot dissociate the processing of socio-emotional information from threat-related information in general. Yet, the contrast between direct and averted gaze allowed to provide some arguments in favor of threat prioritization. Moreover, our experimental design allowed participants to perform two different detection tasks (emotion or color) on the same two-dimensional visual stimulus, while equalizing detection sensitivity across the two tasks. We propose that the prioritization of emotion decisions is driven by the interaction between the social nature of emotions and the behavioral relevance of anger. Still, further work is needed to complement the present findings, by contrasting social but non-threatening stimuli with non-social but threatening stimuli.

Second, although we equalized emotion and color decisions in terms of available sensory evidence and decision difficulty, reflected by similar detection sensitivity parameters, a difference in the reaction times and bias parameters was observed between the two types of decisions. To address this, we used single-trial regressions of EEG signals against the intensity of the displayed emotion and of the color mask, an experimental approach that measures stimulus sensitivity across tasks and which should dissociate perceptual and response bias. We also controlled all brain analyses on a trial-by-trial basis for the participants’ subsequent detection reports (detection of violet in the color task, detection of anger in the emotion task). However, recent evidence in mice suggests that a lack of change in sensitivity does not prevent effects on sensory encoding – and that a behavioral bias can reveal such an effect ^99^. Therefore, we cannot completely exclude that the observed privileged processing in the early time window (100-200ms), which only benefited the emotional, threat-signaling dimension during emotion decisions, may reflect a better sensory encoding of emotion compared to color.

In conclusion, we showed that the emotional, threat-related dimension of our complex two-dimensional stimuli was not only processed, but also translated into action selection earlier than the color dimension. Our findings suggest the existence of prioritized neural computations for processing behaviorally and socially relevant signals, i.e., threat-related facial expressions, during perceptual decision-making. Earlier choice predictive activity over motor cortices during emotion decisions supports the idea that social threat displays are motivationally relevant, not only by providing information about others’ affective states and potential behavioral intentions, but also by conveying action demands to the perceiver ^100^.

## Author contributions

Conceptualization: M.E.Z., V.W., and J.G.; Methodology: M.E.Z., V.W.; Validation: M.E.Z.; Formal Analysis: M.E.Z., R.M., M.S.; Investigation: M.E.Z., E.M.; Data Curation: M.E.Z., R.M., V.W., and J.G.; Writing – Original Draft: M.E.Z., E.M., and J.G.; Writing – Review & Editing: M.E.Z., R.M., M.S., V.W., and J.G.; Visualization: M.E.Z., M.S., and J.G.; Supervision: V.W., and J.G.; Project Administration: V.W., and J.G.; Funding Acquisition: J.G.

## Acknowledgments

This work was supported by FRM Team DEQ20160334878; Fondation de France 00100076; INSERM; ENS and the French National Research Agency under Grants ANR-20-CE28-0003; ANR-10- IDEX-0001-02 and ANR-17-EURE-0017 FrontCog.

## Data and code availability

The datasets generated and/or analysed during the current study, as well as the analyses scripts, will be available in the OSF repository.

