## Supplementary Material for "Prioritized neural processing of social threats during perceptual decision-making"

#equal contribution

### DDM model fitting, selection and reliability check

#### Model fitting

In all experiments Bayesian Hierarchical Drift Diffusion Models have been fitted using the Python package HDDM [1]. The package performs parameter estimates using Markov Chain Monte Carlo (MCMC) sampling. For each model three chains were run in parallel with 5000 iterations each (1500 of which as burn-in period). We used the population-informed priors by Matzke and Wagenmakers (2009)[2] to fit the models.

#### Model Selection

We compared several models by sequentially allowing the drift rate, the boundary separation, the non-decision time, combinations of these or none of them to vary across conditions. We favored the model presenting the lowest DIC (Deviance Information Criterion) and BPIC (Bayesian Predictive Information Criterion [3]) indexes. Al criteria favored as best fitting the model fitting the effect of morph Intensity and Task on the drift rate (v), the threshold (a) and the non-decision time (t0), as well as the Task effect on the starting point bias (z; Table 1).

**Table 1:** DDM model selection results

| model | DIC | BPIC |
|---|---|---|
| v Int Task – a Int Task – t0 Int Task – z Task | 7362.623 | 7803.135 |
| v Int Task – a Int Task – t Int Task | 7469.554 | 789.035 |
| v Int Task – a Int Task – z Task | 10045.345 | 1040.042 |
| v Int Task – t0 Int Task – z Task | 8126.372 | 8478.651 |
| a Int Task – t0 Int Task – z Task | 28373.258 | 28686.442 |
| a Int Task – z Task | 31215.383 | 31447.180 |
| v Int Task – z Task | 12166.646 | 12442.099 |
| t0 Int Task – z Task | 28964.060 | 2920.953 |

Note: Int: Intensity. a: Boundary separation. v: Drift rate. t0: Non-decision time. DIC: Deviance Information Criterion. BPIC: Bayesian Predictive Information Criterion. Bold: best fitting model.

#### Model Diagnostic

Chain Convergence has been visually inspected and verified. All chains presented a 'hairy caterpillar' shape. In Table 2, we provide R-hat and Effective Sample Size (ESS) statistics. An R-hat lower than 1.1 and an ESS in the order of thousands suggest that chains properly converged and mixed 4.

**Table 2:** Mu and Standard Deviation parameter estimates from the best fitting model

| *parameter* | *Estimate* | *95%CrI-low* | *99%CrI-low* | $p_{>0}$ | $p_{<0}$ | *R-hat* | *ESS* |
|---|---|---|---|---|---|---|---|
| *Mu parameters* | | | | | | | |
| *a_Color* | *1.347* | *[1.286, 1.416]* | *[1.263, 1.441]* | *1.000* | *.000* | *1* | *6085* |
| *a_Emo* | *1.340* | *[1.281, 1.403]* | *[1.261, 1.424]* | *1.000* | *.000* | *1* | *8331* |
| *a_Intensity_Color* | *-.068* | *[-.080, -.056]* | *[-.085, -.052]* | *.000* | *1.000* | *1* | *2631* |
| *a_Intensity_Emo* | *-.023* | *[-.034, -.012]* | *[-.038, -.007]* | *.000* | *1.000* | *1* | *3861* |
| *v_Color* | *-.987* | *[-1.194, -.778]* | *[-1.263, -.715]* | *.000* | *1.000* | *1* | *8401* |
| *v_Emo* | *-.211* | *[-.370, -.048]* | *[-.431, .004]* | *.006* | *.994* | *1* | *7922* |
| *v_Intensity_Color* | *.804* | *[.731, .876]* | *[.709, .895]* | *1.000* | *.000* | *1* | *8638* |
| *v_Intensity_Emo* | *.619* | *[.565, .672]* | *[.547, .690]* | *1.000* | *.000* | *1.001* | *9216* |
| *t_Color* | *.402* | *[.377, .430]* | *[.368, .439]* | *1.000* | *.000* | *1* | *9559* |
| *t_Emo* | *.472* | *[.453, .491]* | *[.447, .498]* | *1.000* | *.000* | *1* | *8877* |
| *t_Intensity_Color* | *.003* | *[.001, .004]* | *[.000, .005]* | *.994* | *.006* | *1* | *2671* |
| *t_Intensity_Emo* | *-.003* | *[-.005, -.001]* | *[-.006, .000]* | *.005* | *.995* | *1.001* | *4058* |
| *z_Color* | *.489* | *[.475, .503]* | *[.471, .508]* | $P_{>.5}$=.058 | $P_{<.5}$=.942 | *1* | *3731* |
| *z_Emo* | *.510* | *[.494, .525]* | *[.490, .530]* | $P_{>.5}$=.896 | $P_{<.5}$=.104 | *1* | *4947* |
| *Standard Deviation parameters* | | | | | | | |
| *SD_a_Color* | *.193* | *[.152, .254]* | *[.141, .279]* | *1* | *0* | *1* | *4012* |
| *SD_a_Emo* | *.181* | *[.144, .237]* | *[.134, .260]* | *1* | *0* | *1* | *6051* |
| *SD_a_Intensity_Color* | *.030* | *[.021, .044]* | *[.018, .049]* | *1* | *0* | *1.001* | *1026* |
| *SD_a_Intensity_Emo* | *.029* | *[.021, .041]* | *[.019, .045]* | *1* | *0* | *1* | *1913* |
| *SD_v_Color* | *.620* | *[.494, .800]* | *[.461, .888]* | *1* | *0* | *1* | *6323* |
| *SD_v_Emo* | *.479* | *[.380, .626]* | *[.355, .683]* | *1* | *0* | *1* | *5506* |
| *SD_v_Intensity_Color* | *.211* | *[.167, .275]* | *[.157, .302]* | *1* | *0* | *1* | *6253* |
| *SD_v_Intensity_Emo* | *.156* | *[.122, .203]* | *[.114, .225]* | *1* | *0* | *1* | *6909* |

| | | | | | | | |
|---|---|---|---|---|---|---|---|
| *SD_t_Color* | *.079* | *[.064, .104]* | *[.060, .115]* | *1* | *0* | *1.001* | *6714* |
| *SD_t_Emo* | *.056* | *[.045, .073]* | *[.042, .080]* | *1* | *0* | *1* | *7423* |
| *SD_t_Intensity_Color* | *.005* | *[.004, .007]* | *[.004, .007]* | *1* | *0* | *1* | *2042* |
| *SD_t_Intensity_Emo* | *.006* | *[.005, .008]* | *[.004, .009]* | *1* | *0* | *1* | *2363* |
| *SD_z_Color* | *.147* | *[.114, .188]* | *[.104, .202]* | *1* | *0* | *1.002* | *3731* |
| *SD_z_Emo* | *.168* | *[.136, .208]* | *[.127, .221]* | *1* | *0* | *1* | *4947* |

Note: a: Threshold. t0: Non-decision time. v: Drift rate. Estimate: median posterior estimate. 95% and 99% CrI: 95% and 99% Credible Intervals. $p_{>0}$ and $p_{<0}$: proportion of sampled parameters greater and lower than zero. R-hat: R-hat potential scale reduction factor. ESS: Effective Sample Size.

#### Posterior Predictive Check

In order to assess the reliability of our model we simulated data from the best fitting parameters and we assessed their resemblance to the observed data. Simulated and Observed data presented high correlations for Angry/Violet response proportion and median RT for Angry/Violet and Neutral/Grey responses (Figure 1).

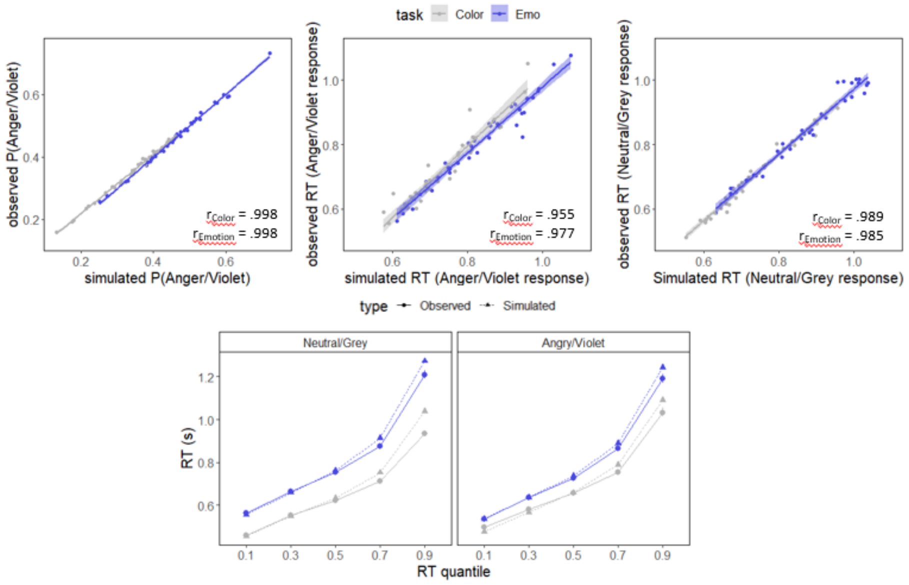


**Figure 1:** Posterior predictive check for the best fitting DDM. Top row: correlation between observed and simulated proportion of avoidance Anger/Violet responses, median RTs for Anger/Violet responses and median RTs for Neutral/Grey responses. Observed and Simulated RTs by quantile of the RT distribution.

#### Parameter Recovery

In order to further assess the reliability of the model we conducted a parameter recovery study by simulating a dataset from the best fitting model and by refitting the same model on this simulated dataset. In this way, we could assess the similarity between the observed and the recovered parameter estimates at both the group- and the individual-level.

All the group-level parameters were successfully recovered, with all the median posterior estimates of the recovered parameters falling within the 95%CrI of the observed parameters (see Table 3, Figure 2). Furthermore, all the individual-level recovered parameters presented high correlations with the observed parameters, beside the effect of Intensity for both the Color and the Emotion task presenting lower correlations ($r_{int\text{-}col}$ = .652, $r_{int\text{-}emo}$ = .695; Figure 3). However, since we based inference on group level parameters, this lower correlation is less of interest.

**Table 3:** Observed and recovered group-level mu parameter estimates

| | Observed parameters | | | Recovered parameters | | |
|---|---|---|---|---|---|---|
| parameter | Estimate | 95%CrI | 99%CrI | Estimate | 95%CrI | 99%CrI |
| a_Color | 1.347 | [1.286, 1.416] | [1.263, 1.441] | 1.367 | [1.304, 1.433] | [1.286, 1.459] |
| a_Intensity_Color | -.068 | [-.080, -.56] ] | [-.085, -.052] | -.067 | [-.081, -.053] | [-.086, -.049] |
| a_Emo | 1.340 | [1.281, 1.403] | [1.261, 1.424] | 1.355 | 1.302, 1.413] | [1.283, 1.436] |
| a_Intensity_Emo | -.023 | [-.034, -.012] | [-.038, -.007] | -.026 | -[.035, -.016] | [-.038, -.013] |
| t_Color | .402 | [.377, .430] | [.368, .439] | .404 | [.377, .432] | [.369, .442] |
| t_Intensity_Color | .003 | [.001, .004] | [.000, .005] | .003 | [.001, .004] | [.001, .005] |
| t_Emo | .472 | [.453, .491] | [.447, .498] | .475 | [.456, .494] | [.450, .501] |
| t_Intensity_Emo | -.003 | [-.005, -.001] | [-.006, .000] | -.003 | [-.005, -.001] | [-.006, .000] |
| v_Color | -.987 | [-1.194, -.778] | [-1.263, -.715] | -1.018 | [-1.218, -.811] | [-1.289, -.739] |
| v_Intensity_Color | .804 | [.731, .876] | [.709, .895] | .826 | [.756, .895] | [.734, .920] |
| v_Emo | -.211 | [-.370, -.048] | [-.431, .004] | -.227 | [-.384, -.073] | [-.429, -.014] |
| v_Intensity_Emo | .619 | [.565, .672] | [.547, .690] | .633 | [.580, .688] | [.560, .705] |
| z_Color | .489 | [.475, .503] | [.471, .508] | .494 | [.481, .507] | [.477, .511] |
| z_Emo | .510 | [.494, .525] | [.490, .530] | .507 | [.493, .522] | [.488, .526] |

Note: a: Threshold. T0: Non-decision time. V: Drift rate. Z: Starting Point Bias. Estimate: median posterior estimate. 95% and 99%CrI: 95% and 99% Credible Intervals.

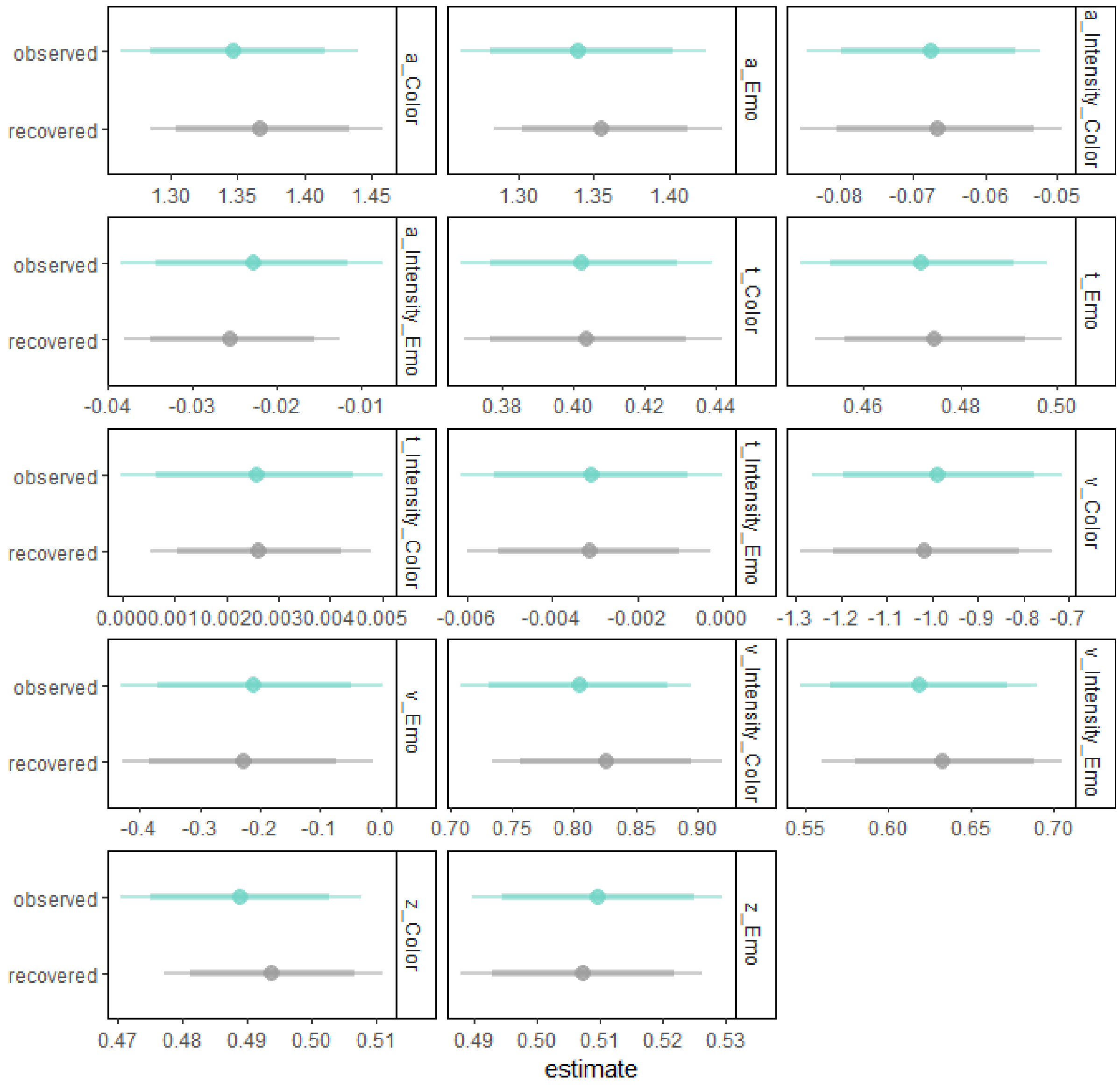


**Figure 2:** Observed and recovered group level DDM parameters. a: Threshold. t0: Non-decision time. v: Drift rate. Z: Starting Point Bias. Points: median posterior estimates. Thick and thin lines 95% and 99% Credible Intervals.

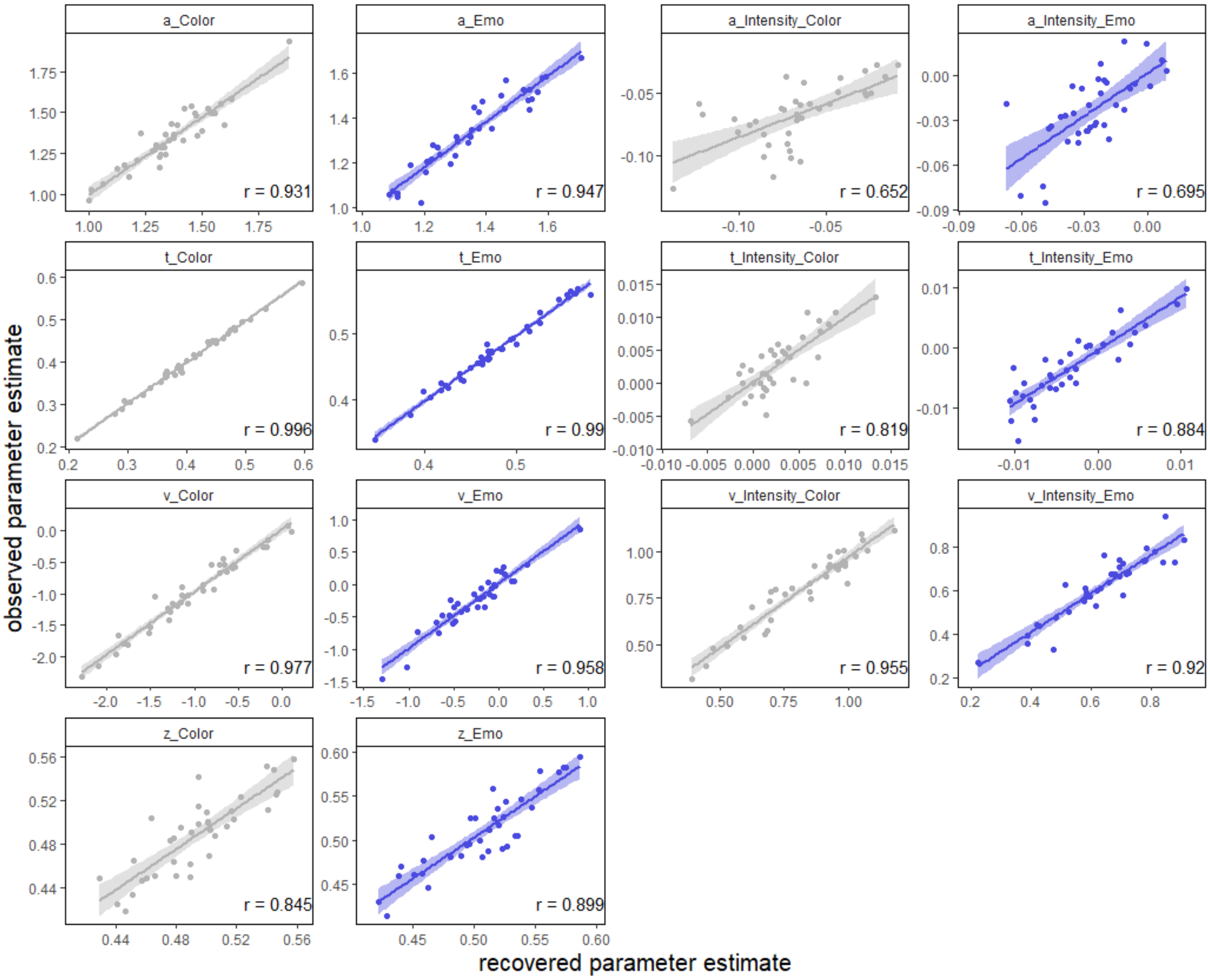


**Figure 3:** Correlations between observed individual DDM parameters and parameters recovered from the simulated dataset. Points: median posterior individual estimates. a: Threshold. t0: Non-decision time. v: Drift rate. z: Starting Point Bias.
